# Primitive GLMY Homology: An Algebraic Topology Approach for the Quantitative Characterization of Graph Pangenomes toward Population Genetic Analysis

**DOI:** 10.64898/2026.07.10.737687

**Authors:** Qi Wu, Jingyan Li, Guoqing Hu, Piyu Zhou, Xin Zhao, Stephen S.-T. Yau

## Abstract

A central task in population genetics is to identify genetic diversity in a population containing a number of individuals. In recent years, with the development of the third generation sequencing (TGS) technology, pan-genome research has become a hot topic. Although graphical representation has been a popular way to represent the pangenome, few works have attempted to describe it in a more mathematical way. In this paper, we used 79 high-quality assembly data of third-generation sequencing in yeast (including*Saccharomyces cerevisiae* and *Saccharomyces paradoxus*) to construct the graph pangenome, and introduced the Primitive GLMY (Grigor’yan-Lin-Muranov-Yau) Homology in algebraic topology to quantitatively represent the pan-genome. We further made an intriguing attempt to conduct a population genetic analysis of this resulting dataset from the topological features of the graph pangenome. We found that there was good agreement between the obtained results and the biological context. We believe this study has developed a method for population genetic analysis of the genetic diversity of genome structural variation.

## Introduction

With the rapid development of high-throughput sequencing technologies and the exponential decrease in sequencing costs, genomics research is gradually transcending reliance on a single linear reference genome, entering the era of the graph pangenome. The concept of the pangenome was initially proposed in bacterial research in 2005, aiming to characterize the collection of all genes and sequence variations within a species, including the core genome shared by all individuals and the variable genome unique to specific individuals (Tet-telin, Masignani, Cieslewicz, et al. 2005). In recent years, pangenome research has moved from proof-of-concept to large-scale application in humans, major crops, and model organisms (Korbel, Urban, Affourtit, et al. 2007; Wang, Mauleon, Hu, et al. 2018; Hufford, Seetharam, Woodhouse, et al. 2021). In particular, breakthroughs based on third-generation long-read sequencing and high-precision assembly technologies have enabled scientists to more completely detect and characterize structural variations, repetitive sequences, and complex alleles that were previously difficult to capture, thereby constructing graph-structured reference genomes capable of encompassing greater population genetic diversity (Liao, Asri, Ebler, et al. 2023). Such graphs intuitively present sequence polymorphism and structural variation paths in the form of nodes and edges (Garrison, Sirén, Novak, et al. 2018), which not only significantly improves the completeness and accuracy of genome alignment and variant detection but also provides unprecedented resolution for deeply resolving species evolutionary history, discovering functional genes, and elucidating the genetic mechanisms of complex traits (Gao, Yang, Chen, et al. 2023). Currently, the graph pangenome is being widely integrated with fields such as precision breeding and disease association analysis, and its construction methods and standardized applications remain one of the most prospective research directions in the field.

Population genetics studies the diversity differences between populations of the same species with different genetic backgrounds and explores the sources and evolution of such differences. Deeply mining the population genetic information contained within pangenome graphs is becoming an increasingly hot research topic. The core of such methods lies in treating every non-core gene or variable sequence block in the pangenome as a genetic trait. By constructing a genome-wide “presence-absence matrix,” classic methods such as Principal Component Analysis (PCA), population structure inference, and phylogenetic reconstruction are applied to reveal population differentiation signals that are difficult to capture via traditional SNP-based analysis (Tettelin, Masignani, Cieslewicz, et al. 2005; Golicz, Bayer, Barker, et al. 2016). In recent years, this approach has achieved a series of breakthroughs in microbial evolution, crop domestication, and human population genetics. On the one hand, new analysis methods and tools are constantly being developed; on the other hand, research in specific species has discovered a large number of structural variations, subspecies, or ecotypes, and even identified various variable genes related to disease resistance and environmental adaptation (Bayer, Golicz, Scheben, et al. 2020). However, this field still faces challenges such as the standardization of computational methods, the accuracy of genotyping complex variant genes, and the integrated analysis of large-scale pangenome data (Garrison, Sirén, Novak, et al. 2018). To deeply resolve the complex relationships and global constraints defined by the directionality of edges (variation paths) within the graph structure, more powerful topological analysis tools need to be introduced.

In recent years, GLMY homology theory has provided a novel perspective for the study of directed networks via algebraic topology. This theory appeared in earlier works under the name path homology and was subsequently developed by Grigor’yan, Lin, Muranov, and Yau in a series of papers published between 2012 and 2015 (Grigor’yan, Lin, Muranov, et al. 2012; Grigor’yan, Lin, Muranov, et al. 2014; Grigor’yan, Lin, Muranov, et al. 2015). For this reason, it is now often referred to as GLMY homology. In this paper, we use the term GLMY homology uniformly throughout.

Its core innovation lies in constructing chain complexes directly based on directed graphs and establishing homology groups by defining *∂*-invariant subspaces, thereby capturing directed flows, cycles, and high-dimensional voids in the network without simplifying directional information into undirected structures. Subsequent studies further distinguished between Regular GLMY Homology and Primitive GLMY Homology, the latter being more advantageous in practical applications. The theoretical basis adopted in this paper is the Primitive GLMY Homology, which was developed and formally established by Li, Muranov, Wu, and Yau based on the aforementioned framework (Li, Muranov, Wu, et al. 2025). Applying GLMY homology to graph pangenome analysis holds the promise of transcending the traditional node-based analysis paradigm. It can detect connected components and directed cycle structures in the directed subgraph formed by the genome sequences of the sampled isolates and the reference isolate within the pangenome. This provides a powerful mathematical framework for revealing population differentiation at the topological level, identifying “topotypes” representing different evolutionary paths, and understanding the impact of complex structural variations on overall genome plasticity.

In this paper, we take isolates of *Saccharomyces cerevisiae* and *Saccharomyces paradoxus* with genome assemblies obtained via third-generation DNA sequencing as an example. We constructed a draft yeast graph pangenome framework. Based on this draft, we introduced primitive GLMY homology to establish a pipeline for the population genetic analysis of the topological structure of the graph pangenome. Specifically, the entire graph was decomposed into several subgraphs, and a method for the quantitative description of each orthologous subgraph was established based on GLMY homology. Consequently, we can treat an orthologous subgraph as the counterpart of a genetic locus; the path of a sampled isolate and the reference isolate on the subgraph form a directed graph, and its topological Betti numbers can be calculated. These topological Betti numbers serve as the allelic state of the given sampled isolate at such a “genetic locus.” This results in a graph pangenome matrix distinct from the node presence-absence matrix. We utilized this matrix to conduct conventional population genomic analyses, including phylogenetic tree construction, population structure analysis, and principal component analysis. Our results show that this method yields results with excellent consistency with biological intuition. This opens up new possibilities for population genetic studies of genomic structural variation.

## Results

### Graph pan-genome

We utilized high-quality genome assemblies of 79 yeast isolates from *Saccharomyces* genus based on thirdgeneration sequencing data to establish a minigraph-based pangenome framework map. The map includes the reference isolate S288C and 63 other isolates from *Saccharomyces cerevisiae* and 15 isolates from *Saccharomyces paradoxus*. Only variations larger than 50bp were retained in the framework graph. In the gfa file, we obtained 29,130 nodes and 30,340 edges. Additionally, we identified 851 specific nodes in the S288C reference isolate and a total of 87,404 isolate-specific nodes in the other 78 isolates. Ultimately, a total of 211,407 edges were obtained. We performed a topology-based diversity analysis on this basic graph pangenome framework.

### The orthologous subgraph for topological-population analyses

We identified nodes conserved across all 79 isolates (referred to as orthologous nodes), totaling 2,056. From these, we further obtained 424 orthologous node pairs that maintained a certain distance, thereby allowing for more complex inter-nodal graphical topology. Since these orthologous node pairs exist in all isolates, they can be used to “cut” a relatively independent subgraph from the entire pangenome graph; we term this an “orthologous subgraph.” One orthologous subgraph represents one given chromosomal sequence segment and all variants in the corresponding segment on the genome of the isolate in the sample.

Each subgraph contains 78 Reference Paired Subgraphs (RPS), corresponding to the 78 sampled isolates. Each RPS is a directed graph composed of the two chromosomal “routes” of the sampled isolate and the reference isolate S288C on the pangenome, starting from the start node and proceeding to the end node. We use primitive GLMY homology to characterize the topological structure of this subgraph, selecting 5 topological features corresponding to 5 Betti numbers (the meanings of the 5 sets of Betti numbers are detailed in the section of Material and Methods). For each topological feature, its value can be obtained for each RPS. In this way, one subgraph yields 78 numbers which correspond to 78 isolates in the sample. These numbers form a topological feature vector corresponding to the topological feature. For one subgraph, the five types of Betti numbers yield five topological feature vectors.

Thus, we ultimately obtained a topological feature matrix of 78 rows and 2,120 columns (424 × 5 = 2120). Alternatively, this can be divided into five topological feature matrices of 78 rows and 424 columns. These (and their combinations) were used for downstream population genetic analyses.

### Reference paired subgraph (RPS) construction and edge-weight encoding

To transform local structural differences in the graph pangenome into quantitative features suitable for population genetic statistical analysis, we decomposed the global graph into a set of local intervals defined by orthologous node pairs and constructed a *reference paired subgraph* (RPS) within each interval. Specifically, nodes appearing in all isolates exactly once were screened from the full graph as ortho-nodes; after sorting by the coordinate order of the reference isolate (S288C), adjacent (or grouped boundary) orthologous nodes formed an interval boundary pair (*s, t*). In this study, a total of 424 orthologous intervals were obtained.

For any interval (*s, t*), we extracted:

- The path of the reference isolate on the graph from *s* to *t*;
- The path of any sampled isolate *X* among the 78 isolates on the graph from *s* to *t*.

Subsequently, the sampled path was reversed to obtain a set of edges from *t* to *s*, which was then merged with the forward edge set of reference isolate to obtain a closed directed subgraph:

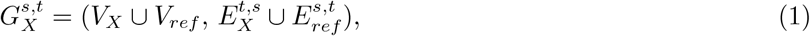

where *E*_*ref*_ is the set of edges for the reference isolate from *s* to *t*, and 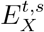 is the set of reversed edges for sampled isolate *X* from *t* to *s*. This closed structure that we termed the reference paired subgraph (RPS) pairs the “interval path of isolate *X*” with the “reference path” into the same topological object, thereby providing the input graph for subsequent GLMY homology calculations.

#### Edge weight encoding (determined by connection symbols)

The path file records directed edges (*u* → *v*) and the connection symbols at both ends (“+”/”−”) line by line. We discretely encoded the combination of connection symbols as weights *w* ∈ {0, 1, 2, 3}:

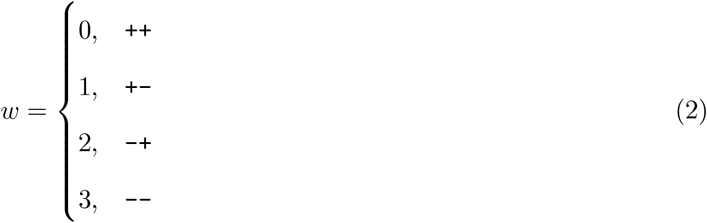

The weights are used as filtration parameters in subsequent persistent GLMY homology calculations.

#### Duplicate edge handling

The weights of edges in 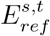 remain exactly the same as in *E*_*ref*_. When merging 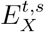 and 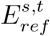, if a co-directional duplicate edge (*u* → *v*) occurs, we retain the record with the smaller weight (i.e., take min *w*) to ensure a deterministic weighted directed graph representation.

### extraction of the topological features

For each reference paired subgraph 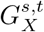 constructed from an orthologous interval (*s, t*), we performed filtration based on edge weights to obtain a sequence of nested weighted directed subgraphs:

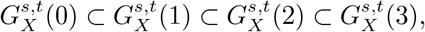

where 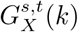 contains all edges with weight *w* ≤ *k*. This filtration process reflects the topological structural relationship between the sampled path and the reference path under different directional consistency thresholds.

On this basis, we calculated the persistent primitive GLMY homology for each 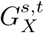 and extracted the following five Betti-related features from the output:

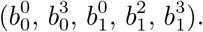

These quantities are not abstract homology dimension indices but are used to characterize *the topological differences of the path structure of isolate X within interval* (*s, t*) *relative to the reference path*. Their specific meanings are as follows:

- 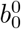: Number of vertices (subgraph scale) which is defined as the number of nodes |*V*_*X*_| in the subgraph 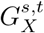. This value reflects the number of distinct graph nodes involved after merging the isolate path and the reference path within the interval (*s, t*). It can be understood as a direct measure of the structural complexity or node expansion of the interval.
- 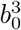: the dimension of 0-dimensional primitive GLMY homology in the final filtration layer 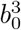 is taken from the 0-dimensional primitive GLMY homology of the maximum filtration layer (i.e., 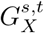(3)). This quantity characterizes the number of connected components in the subgraph after considering all direction combinations. Biologically, it reflects whether the sampled path and the reference path form multiple mutually separated directed structural units.
- 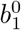: the dimension of 1-dimensional primitive GLMY homology in the low-weight filtration layer 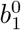 corresponds to the 1-dimensional primitive GLMY homology under a lower weight threshold (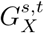 (1)). It mainly characterizes whether non-trivial directed closed structures have formed under conditions of high directional consistency (e.g., “++” or “+-”). This feature is particularly sensitive to detecting cases that are highly consistent with the reference path but contain local rearrangements.
- 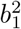: the dimension of 1-dimensional primitive GLMY homology in the medium-weight filtration layer 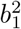 comes from the medium weight threshold (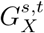 (2)), characterizing the number of directed cycles or non-trivial path closure structures formed when more directional inconsistencies (e.g., “-+”) are allowed. This indicator captures the cumulative effects of structural variations such as inversions and direction flips on the interval scale.
- 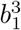: the dimension of 1-dimensional primitive GLMY homology in the final filtration layer 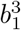 is taken from the maximum filtration layer (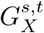(3)), representing the complexity of directed cycle structures contained in the subgraph as a whole after allowing all direction combinations. It comprehensively reflects all local structural differences of the sampled path relative to the reference path and is an overall topological response to insertions, deletions, inversions, and complex rearrangements.

The above five quantities collectively constitute the topological feature representation of each “isolate *X* within orthologous interval (*s, t*)”. After aggregating all 424 orthologous intervals and 78 isolates, a feature matrix is formed for PCA, population structure analysis, and phylogenetic inference. In previous graph pangenome studies, some works directly constructed matrices based on the presence or absence of nodes in the graph for a given isolate and used this “graph-node” matrix for population genetic analysis. To compare with such works, we similarly constructed a node matrix for each subgraph. A straightforward but currently unreported idea is to construct a “graph-edge” matrix using information on the presence or absence of edges in the graph. In this study, we investigated this idea by constructing an edge matrix for each subgraph. In the population genetic analysis, we examined the differences in analysis results between the topological features represented by such node/edge presence-absence information and the topological features defined by our 5 quantities.

### population analyses on the topo-pop dataset

In the 78 × 424 topological feature matrix obtained for each Betti number, every element is a non-negative integer. This is determined by the mathematical properties of homology groups. Thus, this topological feature matrix can be analogized to a microsatellite population genetic sample data table with 78 individuals and 424 microsatellite loci, allowing for various population genetic analyses.

#### PCA results

Based on the 5 topological features extracted from the aforementioned pangenome graph model, we performed PCA clustering analysis on all 78 isolates. The PCA classification results are shown in Figure 3. Generally, in PCA analysis, we hope to both clearly distinguish major biological groups and effectively display differences between different individuals within groups. It can be seen that both 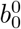 and 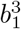 can balance these two requirements, yielding relatively ideal results (Fig. 3A, 3E). Although 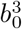 can distinguish the two yeast species well, it does not effectively identify differences between different isolates within *Saccharomyces cerevisiae* (Fig. 3B). Conversely, 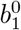 and 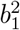 indicated excessive differences between isolates within the species (Fig. 3C, 3D). Simultaneously, the feature combination of 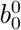 and 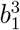 also yielded excellent results. In comparison, results obtained by solely considering node presence/absence information fell between the results of 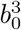 and those of 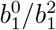. The results obtained by solely considering edge presence/absence information were relatively close to the results of 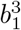.

**Figure 1.**
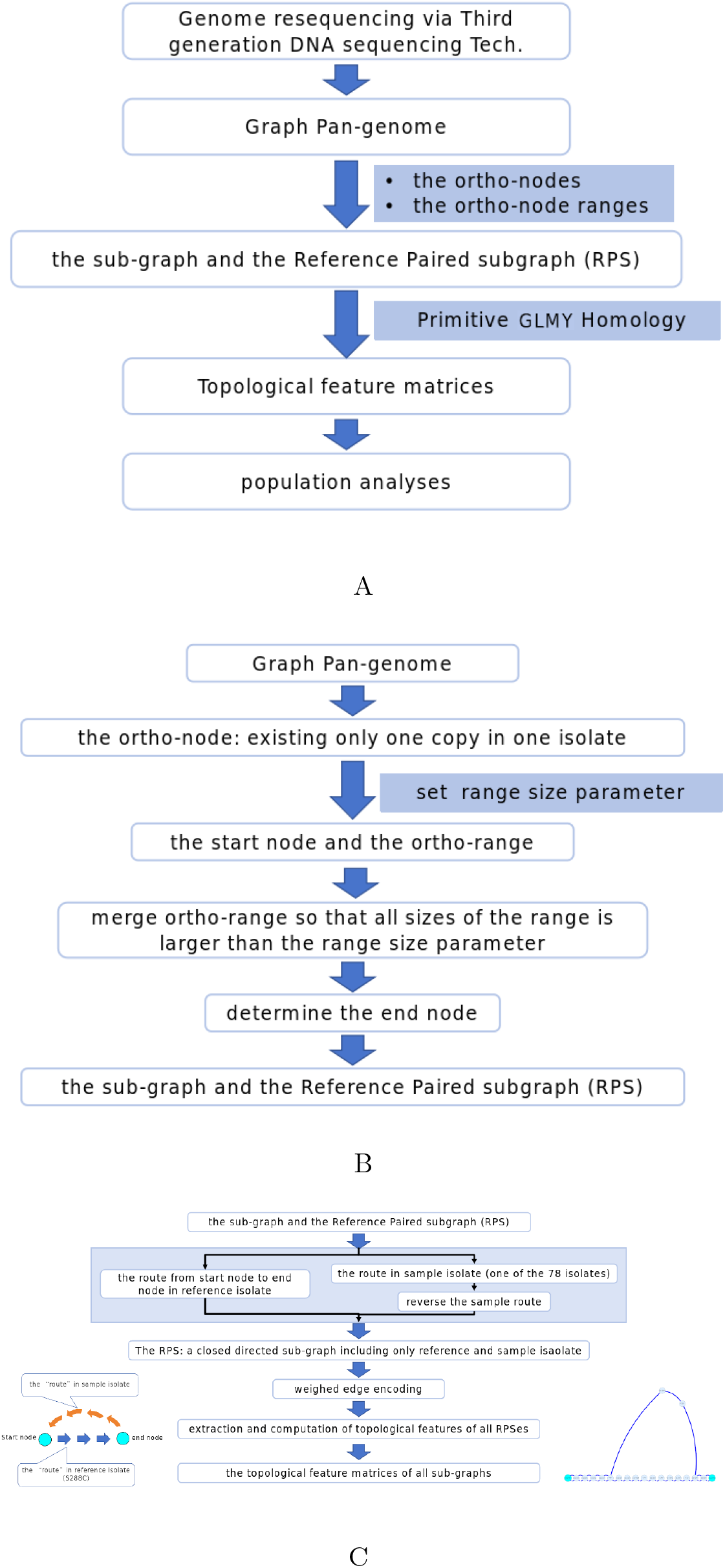
The pipeline of this work: from graph pangenome to population genetic analyses. **A**. overview of the whole pipeline. **B**. the detailed pipeline to obtain the sub-graph. **C**. The detailed pipeline for topological computation. The left-bottom small picture showed the pattern diagram of an RPS, the right-bottom small picture showed a real example of RPS with the ortho-range from start node of s2249 to the end node of s2270 in genome of isolate of GCA_903819185.2.

**Figure 2.**
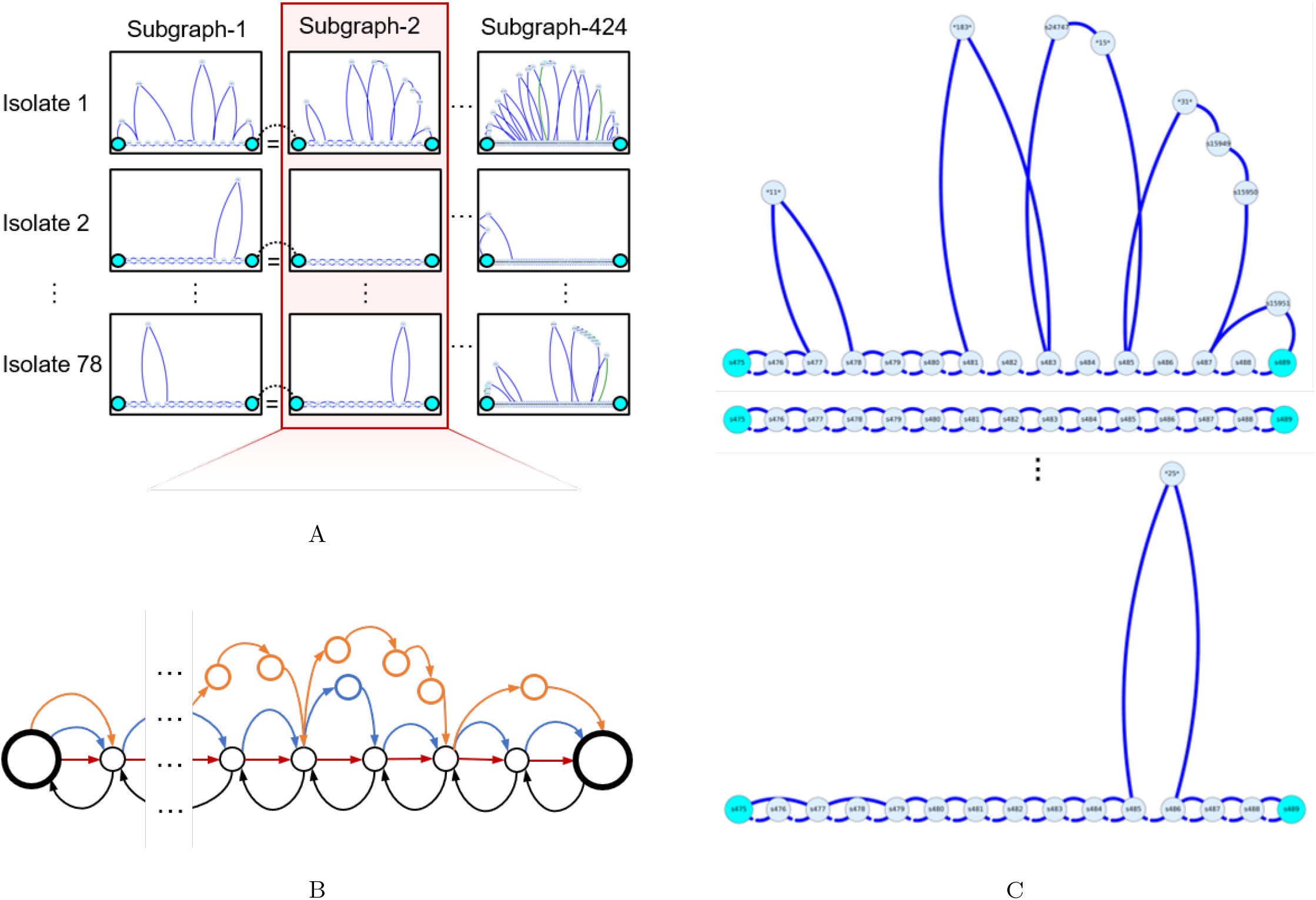
structure of the pangenome graph, subgraph and reference paired graph. **A**. The global view of the total graph. **B**. one example of the RPG with the start and end orthonode of s457 to s489. **C**. Schematic graph of one subgraphs, only 3 RPGs from 3 isolates and the reference isolate (S288C) were shown. In Fig3C, Isolate 1: GCA_030035175.1, Isolate 2: GCA_903819145.2, Isolate 78: GCA_903819195.2. One can see the middle graph showed that the isolate is just the same as the reference isolate with no variation. The bottom graph showed that there is a one-node difference, while the upper graph showed a more complicated and larger variation.

**Figure 3.**
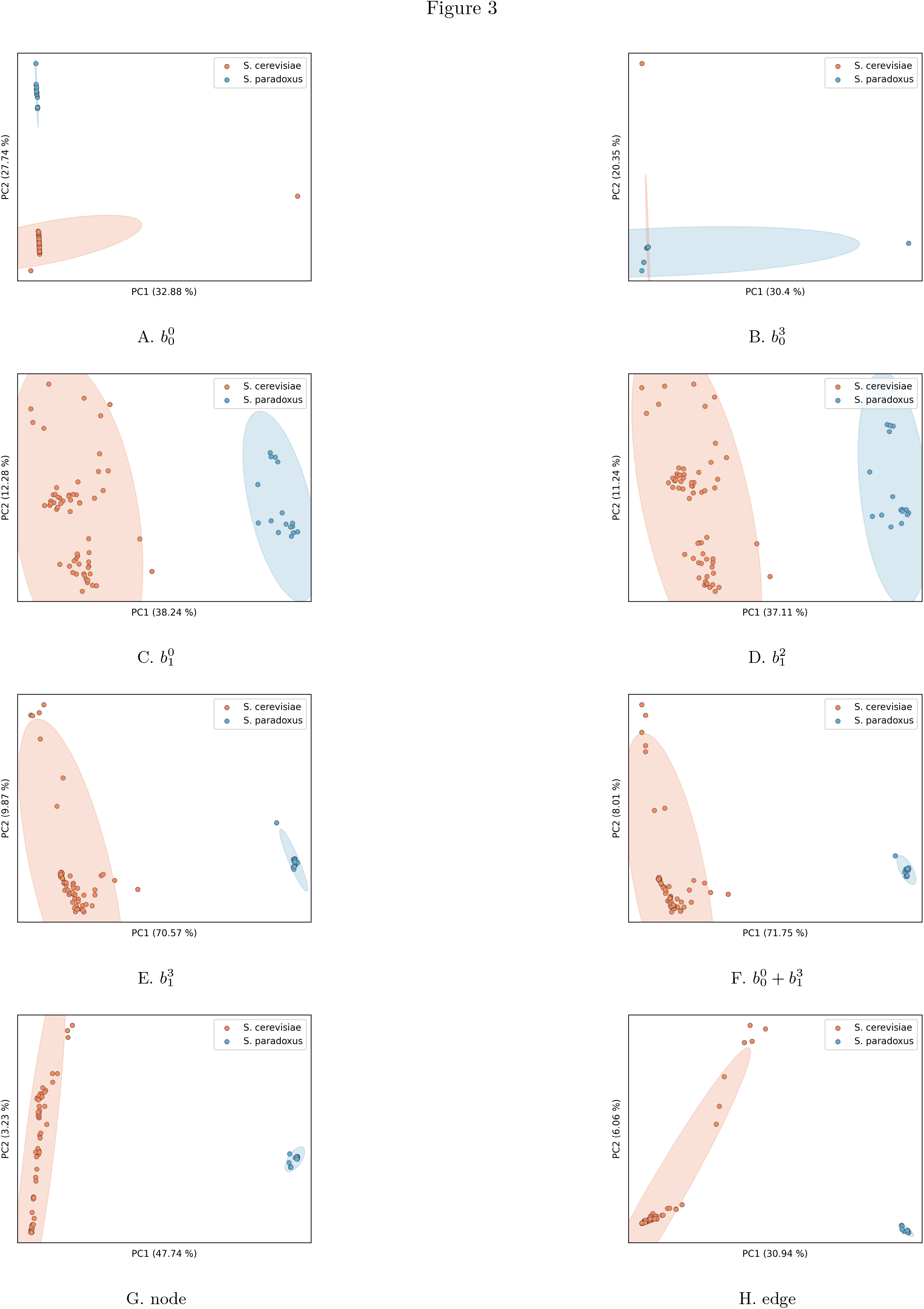
PCA analysis. We used different topological features to perform PCA clustering analysis on the two types of yeast isolates respectively, and marked the 3*σ* confidence ellipses according to the distribution position of each sample in the principal component space. Subfigures a-h correspond to the features used: 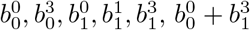, node, and edge.

#### population structure results

Based on the topological features extracted from the aforementioned pangenome graph model, we performed population structure analysis on all 78 isolates. From the perspective of binary classification, the 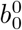 feature and node feature can clearly distinguish *Saccharomyces cerevisiae* and *Saccharomyces paradoxus*. From the perspective of multi-class classification, the node feature remains very stable in dividing into two classes under varying K values. However, this also implies that the “resolution” of the node feature ends here, and it is unable to further distinguish subgroups within the current two yeast populations. The edge feature is also very stable; classification results consistently concentrate into two categories, though some isolates tend to have “uncertain” classification results.

#### Phylogenetic tree results

Based on the topological features extracted from the aforementioned pangenome graph model, we constructed phylogenetic trees for all 78 isolates. The tree construction results are shown in Fig. 4. It can be seen that all three methods can well distinguish the two yeast species, while the internal details of the trees differ slightly. We analyze the details of internal differentiation within *Saccharomyces cerevisiae* later.

**Figure 4.**
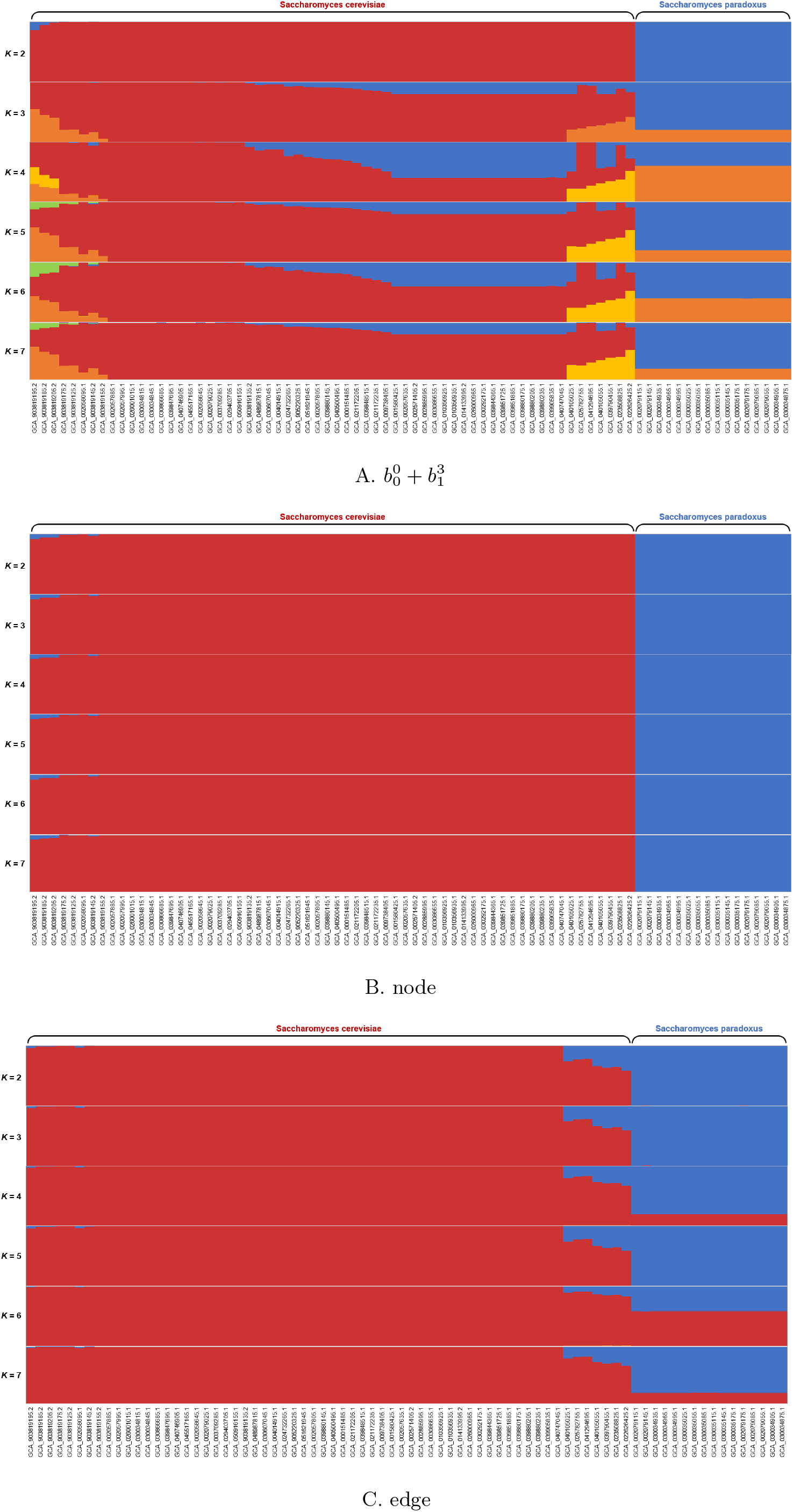
Structure analysis. Subfigures a-c correspond to the population structure analysis results of each isolate under the features 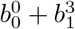, node, and edge, respectively.

**Figure 5.**
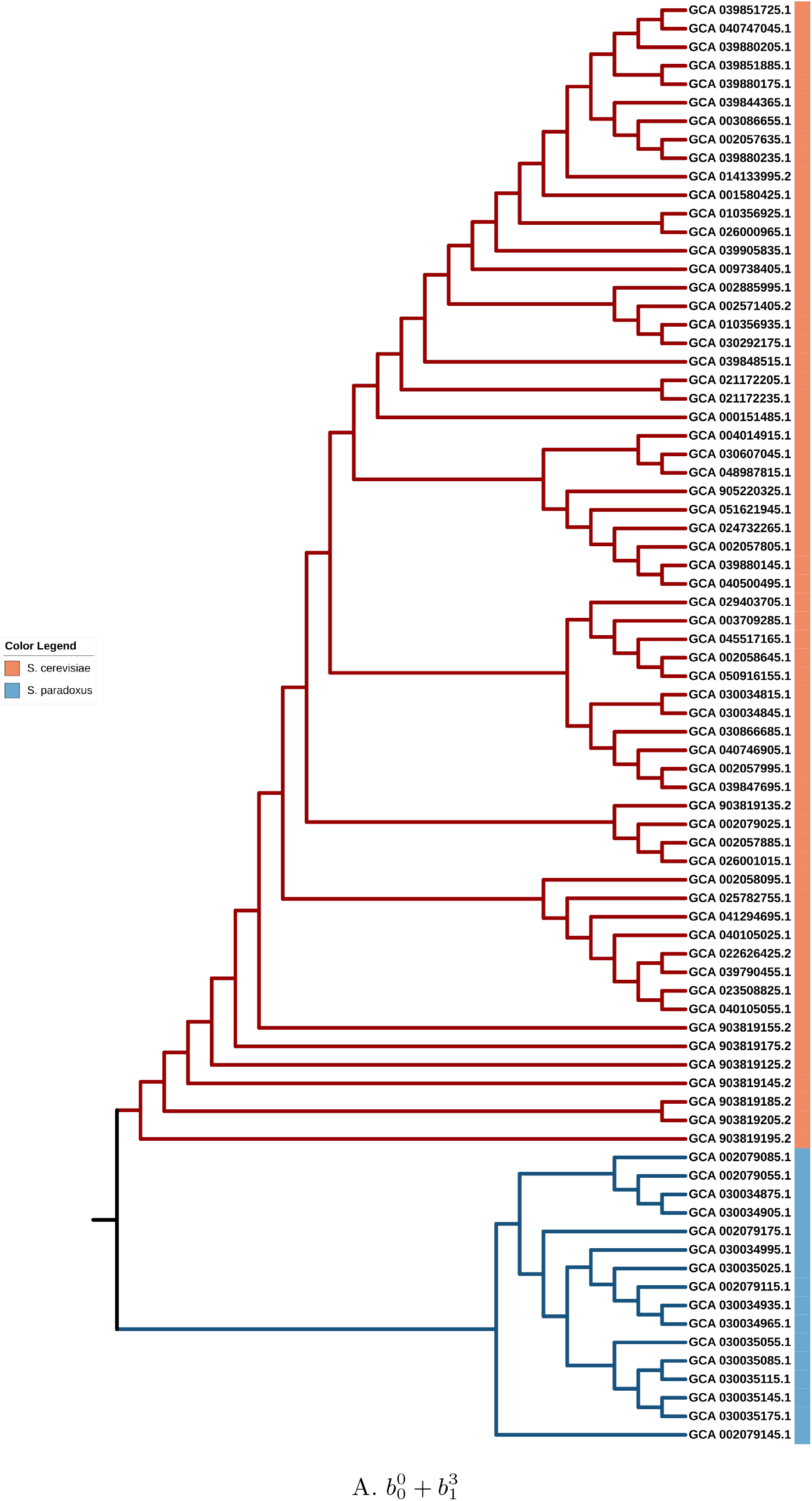
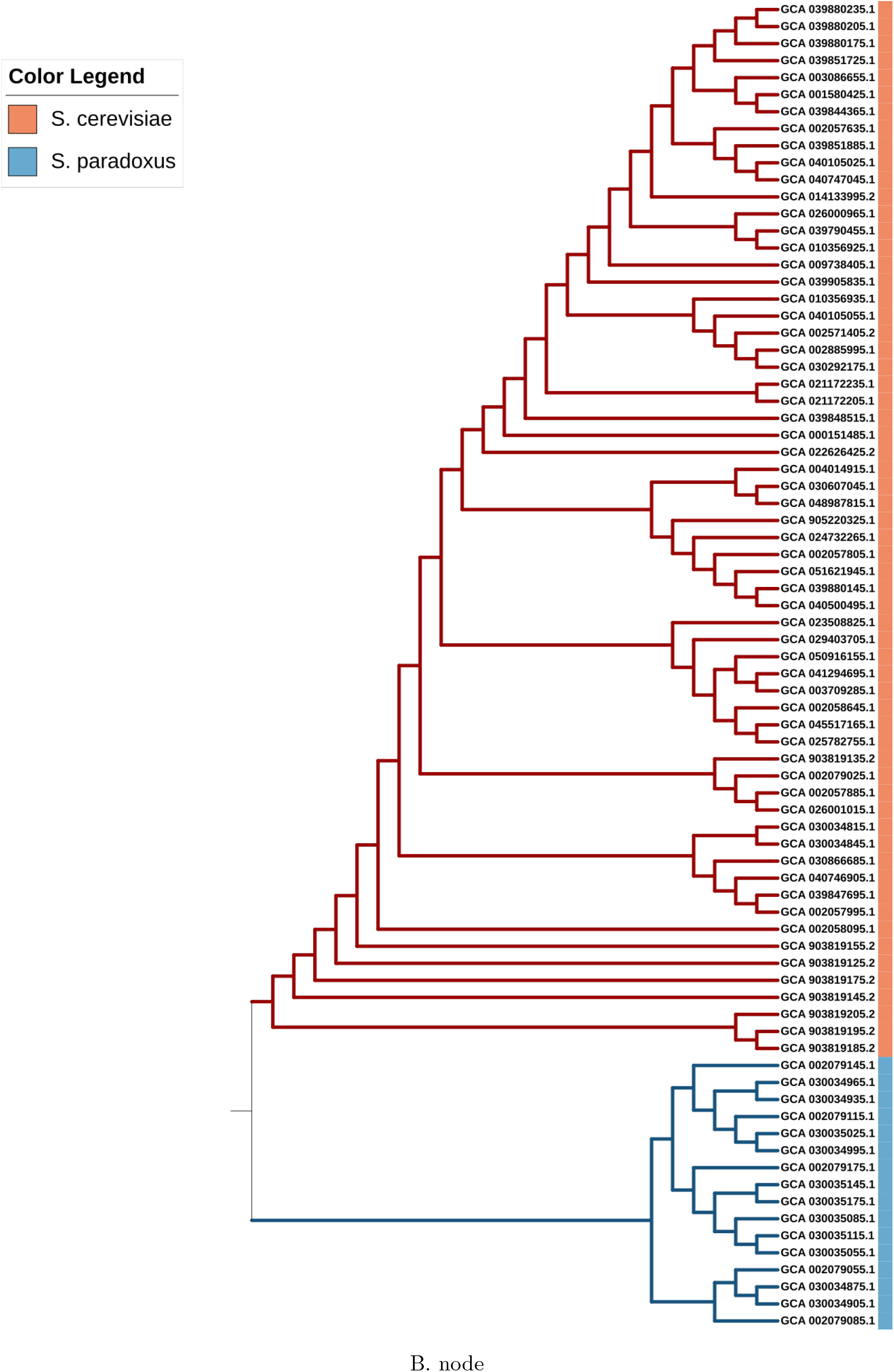
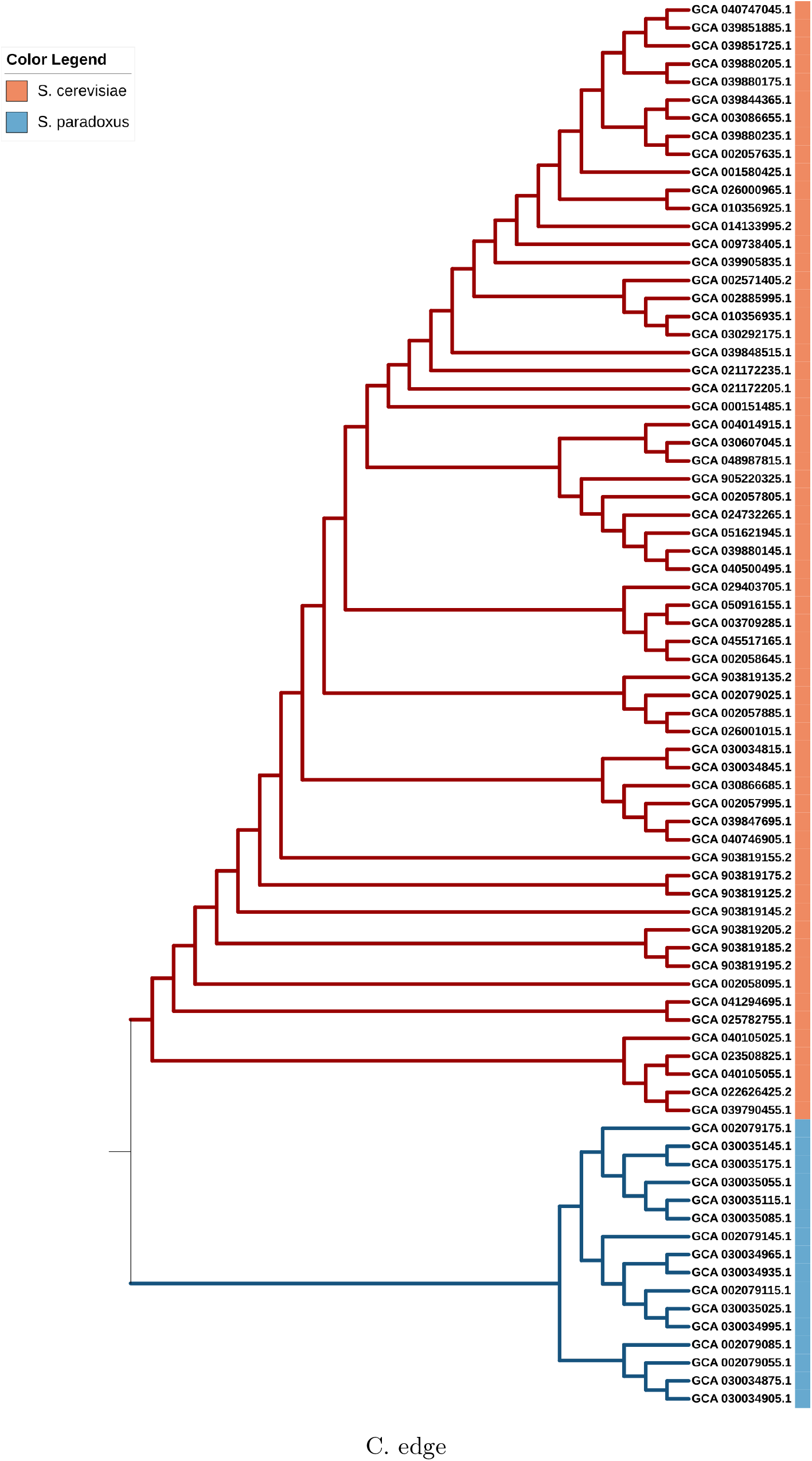
Tree analysis. Subfigures a-c correspond to the phylogenetic tree construction results of each isolate under the features 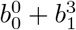, node, and edge, respectively.

#### consistence for two isolate sets in *S. cerevisiae*

From the population genetic structure results, it can be seen that two groups of isolates within *Saccharomyces cerevisiae* exhibit distinct genetic backgrounds. Specifically, in Fig. 4A, when K=3 or a larger value, several isolates on the far left and far right of *Saccharomyces cerevisiae* exhibit unique color combinations. This implies that these isolates have unique genetic backgrounds. We conducted a detailed analysis of these two groups of isolates in the PCA and tree results. These two groups of isolates are:

- isolate set A:GCA_903819195.2, GCA_903819185.2, GCA_903819205.2, GCA_903819175.2, GCA_903819125.2, GCA_002058095.1, GCA_903819145.2, GCA_903819155.2
- isolate set B: GCA_040105025.1, GCA_025782755.1, GCA_041294695.1, GCA_040105055.1, GCA_039790455.1, GCA_023508825.1, GCA_022626425.2

The results are shown in Fig. 6. It can be observed that in the 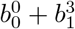 feature combination and the edge presence/absence information results, the positions of these two groups of isolates can be uniquely clustered. However, in the node presence/absence information results, while set X can be well clustered individually, set Y is not clustered and is dispersed throughout the tree.

**Figure 6.**
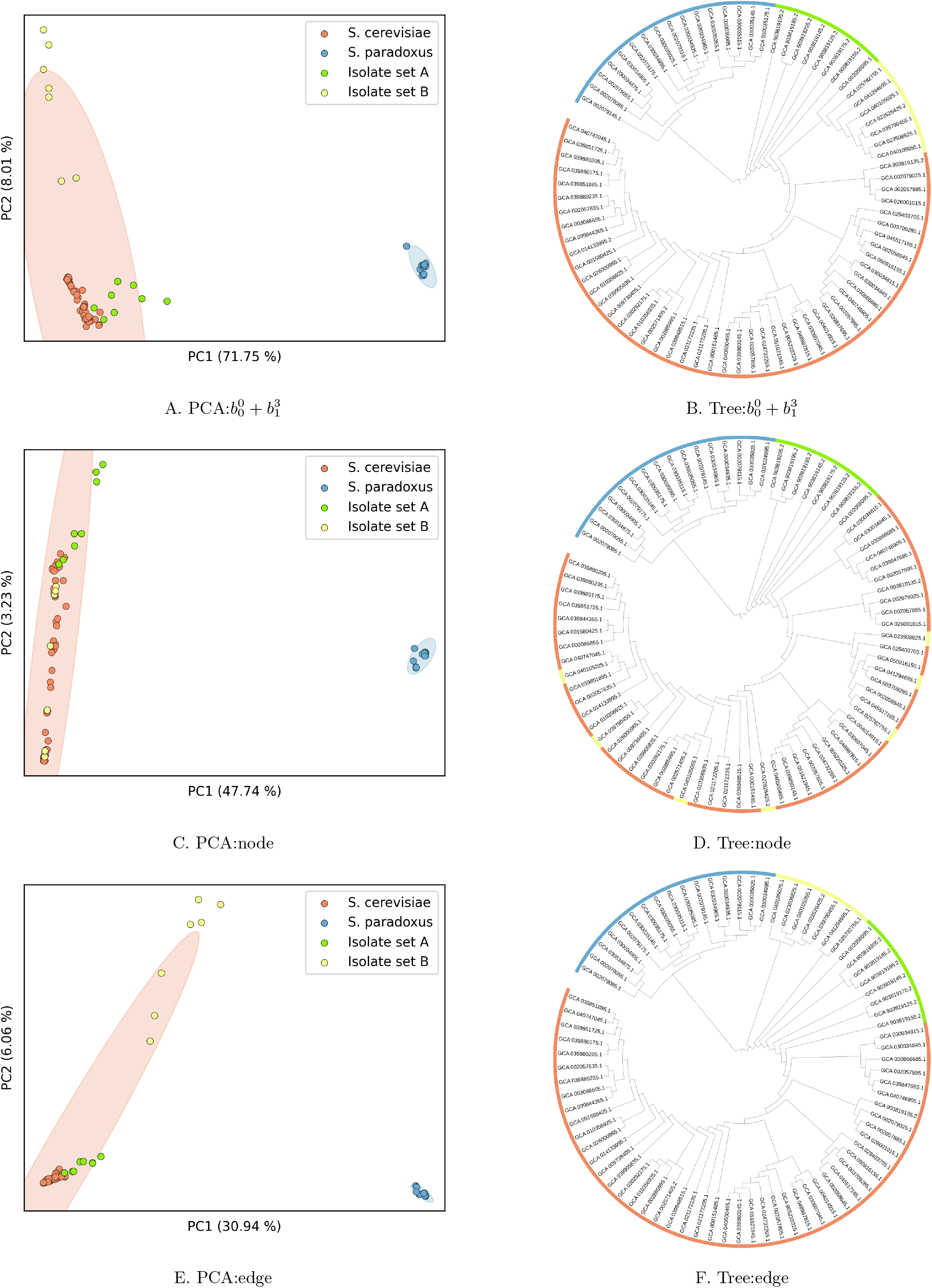
Isolate set analysis. Subfigures a-f correspond to the distance distribution of isolate set A and isolate set B in PCA analysis and phylogenetic tree construction under the features 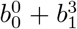, node, and edge, respectively.

## Discussion

This study constructs a complete methodological framework from graph pangenome to population genetic analysis. By decomposing the global graph into subgraphs defined by orthologous nodes and constructing “Reference Paired Subgraphs” (RPS), we transformed the complex problem of comparing genomic structural variations into a series of standardized directed graph topology analysis problems. By introducing Primitive GLMY Homology, an algebraic topology tool, we successfully achieved a rigorous mathematical characterization of RPS topological structures and extracted multi-dimensional Betti number features. Finally, PCA, population structure inference, and phylogenetic analysis based on these topological feature matrices yielded results highly consistent with biological species classification and known isolate genetic backgrounds. This demonstrates for the first time that topological quantification based on GLMY homology can effectively capture population genetic signals inherent in genomic structural variations, opening a completely new and mathematically grounded avenue for graph pangenome analysis.

The core success of our method in population genetic analysis lies in our implementation of a novel biological analogy: treating each orthologous subgraph as a generalized genetic locus, and treating the specific Betti number (a non-negative integer) calculated for an isolate in that subgraph RPS as a generalized allelic state at that locus. Thereby, the entire graph pangenome is transformed into an “individual *X* topological locus” topological feature table. The specific statistical feature values in each cell of the table are transformed into different “topological allelic states” at the “topological locus.” Considering that the variety of statistical feature values for the same topological feature across different isolates is finite and generally greater than 2, the form of this topological feature table is isomorphic to traditional multi-allelic genetic data. Analysis results show that clustering and grouping based on the topological feature matrix are significant, and certain features (such as the 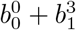 combination) outperform traditional “node/edge presence-absence” matrices in revealing fine internal population structures. This suggests that topological features not only reflect genetic differences but their capture of “higher-order” structural relationships (such as connectivity and cyclic structures) may be closer to evolutionary processes driven by complex variations like inversions and translocations, thus validating the inherent rationality of analogizing topological invariants as novel genetic markers.

It is worth noting that not all topological features perform consistently in population analysis. For example, 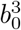 (0-dimensional homology of the final layer), while clearly distinguishing species, is insensitive to intraspecies variation; whereas 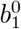 and 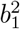 (1-dimensional homology of low/medium weight layers) amplify intra-species differences, potentially leading to over-stratification. The best overall performance, achieved by 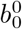 (number of nodes) and 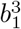 (global 1-dimensional homology), provides information from two complementary dimensions: structural complexity and global cyclic structure, respectively. This differentiation precisely reveals that different topological features map to distinct biological processes: 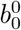 may more directly reflect sequence content changes caused by insertions/deletions, while 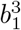 is more sensitive to complex rearrangements (such as local inversions and translocations within chromosomes) requiring global coordination. Therefore, the selection and combination of features should be based on specific biological questions, and the systematic comparison conducted in this study provides a practical basis for this.

To simplify calculations, we have not yet addressed the repetitive occurrence of node sequences and translocation variations between chromosomes, which may miss some important global topological constraints. Future work could incorporate these complex factors by introducing multi-layer graph or hypergraph models. Secondly, the current method uses Betti numbers as summary statistics. In persistent homology, “barcodes” are used to handle sequence directionality information. Based on current results, the analysis results of 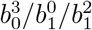 are not significantly better than 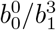, so the advantage of persistent homology in reflecting evolutionary processes does not seem to manifest in sequence directionality. Developing new methods based on persistence diagrams to reflect certain evolutionary relationships should be a focus of future exploration.

Traditional population genetics considers diversity and its evolution in a sample of individuals under the locus-allele framework. This approach cannot account for the diversity of DNA sequence (chromosomal) structural changes and their evolution. The study of topological features of the graph pangenome in this paper pioneers a new line of thought, shifting from focusing on site segregating states under the premise of sequence linearity to focusing on connectivity and shape in sequence space. This new perspective can encompass more general genotypic variation states and provides the possibility to describe larger genotypic differences (such as sequence differences between species or even higher taxonomic units). In the future, by considering more comprehensive topological invariants and introducing more classic population genetic parameters, we hope to establish a “Topological Population Genetics” theory to quantitatively resolve how evolutionary forces such as natural selection and genetic drift shape the evolutionary trajectories of species directly from the perspective of overall genomic structural plasticity. This may provide fresh insights for bridging macro-evolutionary patterns (such as speciation and adaptive radiation) and micro-genetic variations.

## Materials and Methods

### Genome assembly data set in *Saccharomyces cerevisiae*

We downloaded genome assembly results of *Saccharomyces cerevisiae* and *Saccharomyces paradoxus* isolates obtained using third-generation DNA sequencing technology from the NCBI genome database. These data were used to construct the minigraph pangenome graph. The specific list of isolates is provided in Supplementary Table 1.

### Construction of graph pan-genome

We made a pipeline based on software of minigraph (version 0.21-r606) and minimap to construct pangenome variation graph. We first constructed the graph with only SVs larger than 50, including the following steps:

Step 1: The gfa file obtained by processing the original isolate genome fasta sequences using minigraph includes two types of lines: S lines and L lines. S lines start with the character “S”, recording node information including sequence information, position in the reference genome, etc. L lines start with the character “L”, recording edge information, including which two nodes the edge connects, etc. Step 2: We used gfatools (version 0.4-r214-dirty) combined with Python scripts to generate a gfa file with added P lines based on this gfa file. Step 3: A Python script was used to count how many isolates each node appeared in, and the results were appended to the end of the S lines. The field “SC:Z” records the specific isolate IDs where the node appeared. The field “SP:Z” records in how many isolates this node appeared and the total number of occurrences. This final result file is referred to as the minigraphX_P_SCSP.gfa file, where the letter *X* represents the file version number.

### Identification of orthologous subgraph

In the script used to obtain the minigraph_P_SCSP.gfa file above, we simultaneously counted nodes that appeared in all isolates and appeared exactly once. These nodes are termed “orthologous nodes.” They were arranged according to the order on the chromosomes of the yeast reference isolate S288C and then merged into groups of approximately 10 nodes to obtain ortho-node ranges. The merging principles were as follows:

- Use the first ortho-node as the start node.
- Set the range size parameter to 10.
- Find the 10th ortho-node sequentially from the start node.
- If there are more than 10 nodes from this 10th ortho-node to the end of the chromosome, then this 10th node is the end node.
- If there are fewer than 10 nodes from this 10th ortho-node to the end of the chromosome, then these remaining nodes (fewer than 10) are merged into this range, and the last ortho-node on the chromosome serves as the end node.

Each group of orthologous nodes obtained in this way, defined by the start and end nodes, is called an orthologous pair. The interval in between is called the orthologous range, and this part of the graph pangenome within the ortho-range constitutes a subgraph.

For any given pair of ortho-nodes, it exists in every isolate and appears exactly once. Therefore, in any given isolate X, a directed route of the pangenome connecting several nodes can be traced between these two orthologous nodes. Specifically, this is a path connecting several nodes with several directed edges; let us call it path X. Meanwhile, this pair of nodes also appears exactly once in the genome of the reference isolate. Thus, a path can also be traced in the reference genome; let us call it path R. Noting that path X and path R share common start and end points, the two paths of isolate X and reference isolate R on this chromosomal segment can form a loop. That is, starting from the orthologous node serving as the start point, one travels along the pangenome path of isolate X to the end node, and then returns from the end node to the start point along the pangenome path of the reference isolate. For any given isolate, a pangenome loop can always be constructed in combination with the reference isolate using this method. It should be pointed out that between path X and path R, due to the existence of chromosomal structural variations such as inversions, this loop may exhibit various special topological structures such as overlaps, loops, and knots, rather than a simple circle.

Using a similar method, for any other given isolate Y, path Y also exists for the given orthologous node pair. Thus, a loop including path Y and path R can also be established between isolate Y and reference isolate R. Let us call the loop of isolate X and reference isolate R the X-R loop, and the loop of isolate Y and reference isolate R the Y-R loop. An easily observable point is that the difference between the X-R loop and the Y-R loop is mainly caused by the pangenome differences between isolate X and isolate Y within this orthologous node pair. An extreme example is that if the sequences of isolate X and Y on this chromosomal segment are completely identical, then the X-R and Y-R loops would completely coincide. Roughly speaking, the greater the similarity between isolate X and isolate Y, the greater the similarity between the X-R and Y-R loops; conversely, the smaller the similarity between isolates, the smaller the degree of similarity between the loops. This difference in loop topological structure similarity is a manifestation of genetic diversity between isolates, which can be strictly mathematically described using the method of primitive GLMY homology in topology.

### The primitive GLMY homology

We briefly introduce Primitive GLMY Homology (Li, Muranov, Wu, et al. 2025).

Let *G* = (*V, E*) be a directed graph, where *V* denotes the set of vertices and *E* denotes the set of directed edges in the directed graph. An **allowed** *n***-path** in *G* is a sequence of vertices *i*_0_*i*_1_…*i*_*n*_, satisfying that for any adjacent vertices *i*_*k*_ and *i*_*k*+1_, there is a directed edge *i*_*k*_ → *i*_*k*+1_ in *G*. With coefficients in Z_2_ = {0, 1}, the vector space generated by all allowed *n*-paths in *G* as a basis is denoted as A_*n*_(*G*), i.e.,

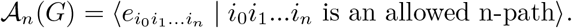

An allowed path *i*_0_*i*_1_…*i*_*n*_ is denoted as 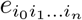 in A_*n*_(*G*), called an elementary allowed path. Elementary allowed paths of length *n* constitute the generators of A_*n*_(*G*).

For each elementary allowed path 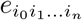 in A_*n*_(*G*), define the boundary operation:

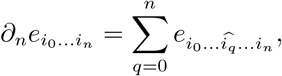

When *n* = 0, define 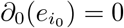. *∂*_*n*_ is called the *n*-th boundary operator.

Generally, when *n* ≥ 2, the result of the boundary operation on an elementary allowed path of length *n* is not necessarily an allowed path. The authors introduced the subspace of A_*n*_(*G*) composed of *∂*-invariant paths:

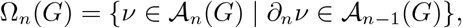

called the subspace of *∂*-invariant paths.

For every path *α* in Ω_*n*+1_(*G*), applying the boundary operator *∂*_*n*+1_ results in an allowed path *∂*_*n*+1_(*α*) of length *n*, i.e., *∂*_*n*+1_(*α*) ∈ A_*n*_(*G*). Furthermore, *∂*_*n*_(*∂*_*n*+1_(*α*)) = 0, i.e. every path in Ω_*n*+1_(*G*) becomes 0 after two boundary operations, this means that the image of *∂*_*n*+1_ is contained in the kernel of *∂*_*n*_ (the subspace composed of all *n*-allowed paths *ν* satisfying *∂*_*n*_(*ν*) = 0). Thus, we can define the quotient space:

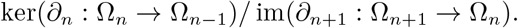

This quotient space is called the *n*-th primitive GLMY homology group of the directed graph *G*, denoted as *H*_*n*_(*G*). Its dimension is denoted as *β*_*n*_, called the *n*-th Betti number of the directed graph *G*.

From the definition of the quotient space, every non-zero element in *H*_*n*_(*G*) is represented by a cycle whose boundary vanishes, and which is not the boundary of any *n* + 1-allowed path. In this paper, we only use 0-dimensional and 1-dimensional primitive GLMY homology. Similar to the homology of simplicial complexes, the 0-dimensional and 1-dimensional primitive GLMY homology of a directed graph can also detect structural information in the directed graph, and these features can be characterized by non-negative integers. For example:

- *β*_0_ detects the number of connected components of the directed graph, i.e., how many mutually unconnected parts the directed graph can be divided into;
- *β*_1_ detects the number of linearly independent closed loops formed by 1-paths that cannot be contracted to a point;

The figure below shows several non-trivial 1-dimensional closed loop shapes that frequently appear in this paper.

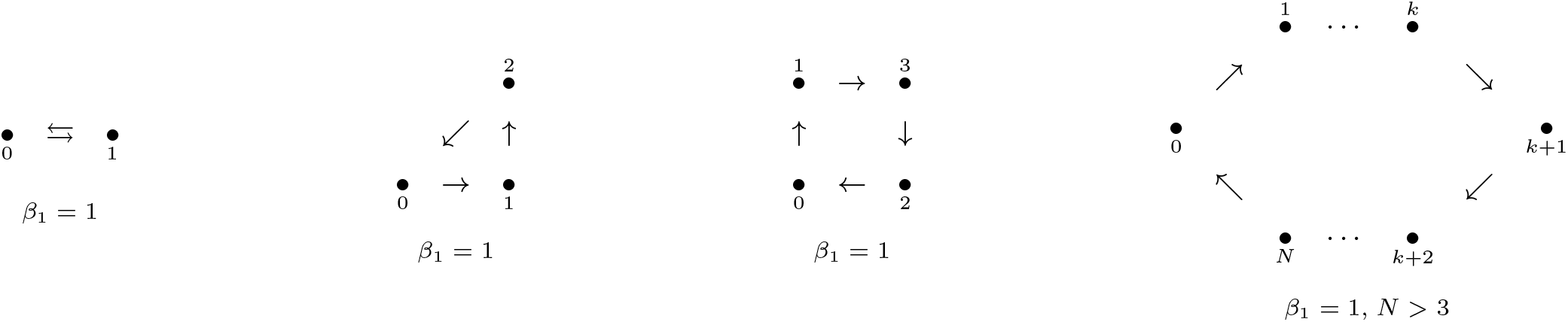

### The persistent primitive GLMY homology

Persistent primitive GLMY homology provides a natural tool for characterizing the evolution of the topological structures of directed graphs across multiple scales. For graphs constructed from biological gene sequences, the graph structure often depends on a chosen scale parameter (e.g., edge weights). By applying a filtration with respect to this parameter, we obtain a nested sequence of directed graphs

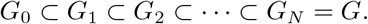

Within this framework, for each homology dimension *n* ≥ 0, the filtration gives rise to a sequence of homology groups

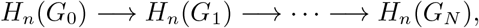

in which each map is induced by the inclusion *G*_*i*_ *‘*→ *G*_*i*+1_.

In this process, the *n*-th Betti number may vary as the filtration scale changes. As the parameter increases, new *n*-dimensional cycles may emerge, leading to an increase in *β*_*n*_, while previously nontrivial cycles may become boundaries in a larger graph, resulting in a decrease in *β*_*n*_.

For each homology dimension *n* ≥ 0, the filtration therefore produces a sequence of Betti numbers

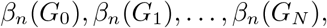

where *β*_*n*_(*G*_*i*_) = dim *H*_*n*_(*G*_*i*_). The scale-dependent behavior of these Betti numbers characterizes the multiscale topological structure of the nested directed graph sequence, providing insight into structural repeatability and modular organization in complex biological systems.

### PCA analysis on the topo-pop dataset

Following the feature analysis in the aforementioned steps, we employed Principal Component Analysis (PCA) to reduce the dimensionality of the high-dimensional topological feature matrix and visualize it. As mentioned in the previous, we selected 5 topological features, as well as the number of nodes and edges in the pangenome graph, totaling 7 features, to perform principal component analysis separately and compare the results. We retained the first two principal components explaining the largest variance and obtained the coordinates of each population’s genomic data on PC1 and PC2, as well as the percentage of variance explained. To assess the stability of clustering, we further calculated the 3*σ* confidence ellipse for each group: using the coordinates of samples on the PC1-PC2 plane as input, we estimated their 2 × 2 covariance matrix, solved for eigenvalues and eigenvectors, and drew ellipses with semi-axis lengths of 3*σ*, thereby intuitively displaying the degree of genetic separation under different topological feature combinations and potential admixture areas.

From the 5 topological features, we selected two features, 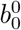 and 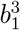, and combined them for PCA analysis. To eliminate differences in dimensional scale, both features were max-normalized.

### Population structure analysis on the topo-pop dataset

Population structure inference was performed with the Bayesian clustering software STRUCTURE v2.3.4. A total of 78 isolates genotyped at feature-corresponding-number binary loci. (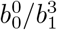 feature for 424 respectively, 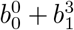 feature for 848, node feature for 79490, and edge feature for 211407) were treated as haploid (ploidy = 1) and supplied in a single-row-per-individual format. Allele coding followed the”-9 = missing”convention; no prior population information, geographic labels, or phenotype data were used (POPDATA = POPFLAG = LOCDATA = PHENOTYPE = 0). For each assumed number of clusters (*K*) from 2 to 7, thirty independent runs were launched to assess stochastic variability among chains. Each run used 10 000 burn-in iterations followed by 10 000 Markov-chain Monte-Carlo (MCMC) replicates. The admixture model with correlated allele frequencies was employed (NOADMIX = 0; FREQSCORR = 1), and the degree of admixture (*α*) was inferred from the data starting at = 1.0 with a uniform prior (UNIFPRIORALPHA = 1; ALPHAMAX = 10; ALPHAPROPSD = 0.025). No linkage model or correction was applied (LINKAGE = 0; LAMBDA = 1). Proposal tuning parameters were set to default (METROFREQ = 10; UPDATEFREQ = 1), and ancestry inference was based solely on genotype likelihoods (PFROMPOPFLAGONLY = 0). Convergence was monitored by visual inspection of log-likelihood traces; all runs reached stationarity within the burn-in phase. The optimal K was evaluated by plotting the log probability of the data [*L*(*K*)] and Δ*K* (Evanno et al. 2005) across runs. Individual ancestry coefficients (Q) from the thirty replicates were aligned with CLUMPP v1.1.2 (Greedy algorithm, 1 000 random input orders) and visualised with DISTRUCT v1.1.

### Phylogenetic tree analysis on the topo-pop dataset

To validate the effectiveness of these several types of topo-feature in biological classification, we constructed phylogenetic trees using (Neighbor-Joining, NJ) method. Phylogenetic reconstruction centered on topological feature distance. First, starting from the 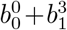 topological feature matrix (78×(424×2) matrix), we calculated the upper triangular distance matrix using Euclidean distance to measure differences between samples. Based on this distance matrix, a neighbor-joining tree was constructed, iteratively finding the minimum evolution tree under unrooted conditions, where branch lengths directly reflect relative differences in feature space. Similarly, we processed the node count feature (78 × 79490 matrix) and edge feature (78 × 211407 matrix) using the same steps. All the above steps were implemented in Python code. The phylogenetic tree structure was visualized using iTOL (v7.4.2, online).

## Supporting information

supplemental Information including Tables and Figures.

Supplemental Tables with the information of the yeast isolates

## Funding

This work is supported by the National Natural Science Foundation of China (No. 12201015 and No. 12171275), and Tsinghua University Education Foundation, and the Beijing Natural Science Foundation (Grant No. IS25032 and the International Scientists Project, Grant No. IS25081).

## Notes

### Competing Interest Statement

The authors have declared no competing interest.

