## supplemental Information including Tables and Figures. for "Primitive GLMY Homology: An Algebraic Topology Approach for the Quantitative Characterization of Graph Pangenomes toward Population Genetic Analysis"

A series of analyses were performed with the result figures shown below.

**Table S1** The information of the isolate and the genome assembly.

**Figure 1** Population structure analysis results on the $b_0^0$ and $b_1^3$ feature for 424 respectively. Panel **A** for the feature of $b_0^0$, and panel **B** for the feature of $b_1^3$. These two results exhibited discriminative abilities comparable to those of the $b_0^0 + b_1^3$, node and edge features, particularly for the $b_1^3$ feature.

**Figure 2** The correlation between the two species of the two slected topological features. Panel **A** for the feature of $b_0^0$, and panel **B** for the feature of $b_1^3$. Each orthologous node showed one point in the figure. The x-axis showed the mean of the values of isolates from *S. paradoxus*, and the y-axis showed those from *S. cerevisiae*. It can be seen that the $b_1^3$ illistrated more diversity between the two yeast species.

**Figure 3** The information of the 424 orthologous nodes along the 16 chromosome in yeast genome. Panel **A** showed the feature of $b_0^0$, and panel **B** for the feature of $b_1^3$. For each panel, the upper subfigure showed the comparison of the two species, the lower subfigrue showed the difference of the two species. It can been seen that for both of the tow features, their values are generally higher in *S. cerevisiae* than those in *S. paradoxus*.

**A**

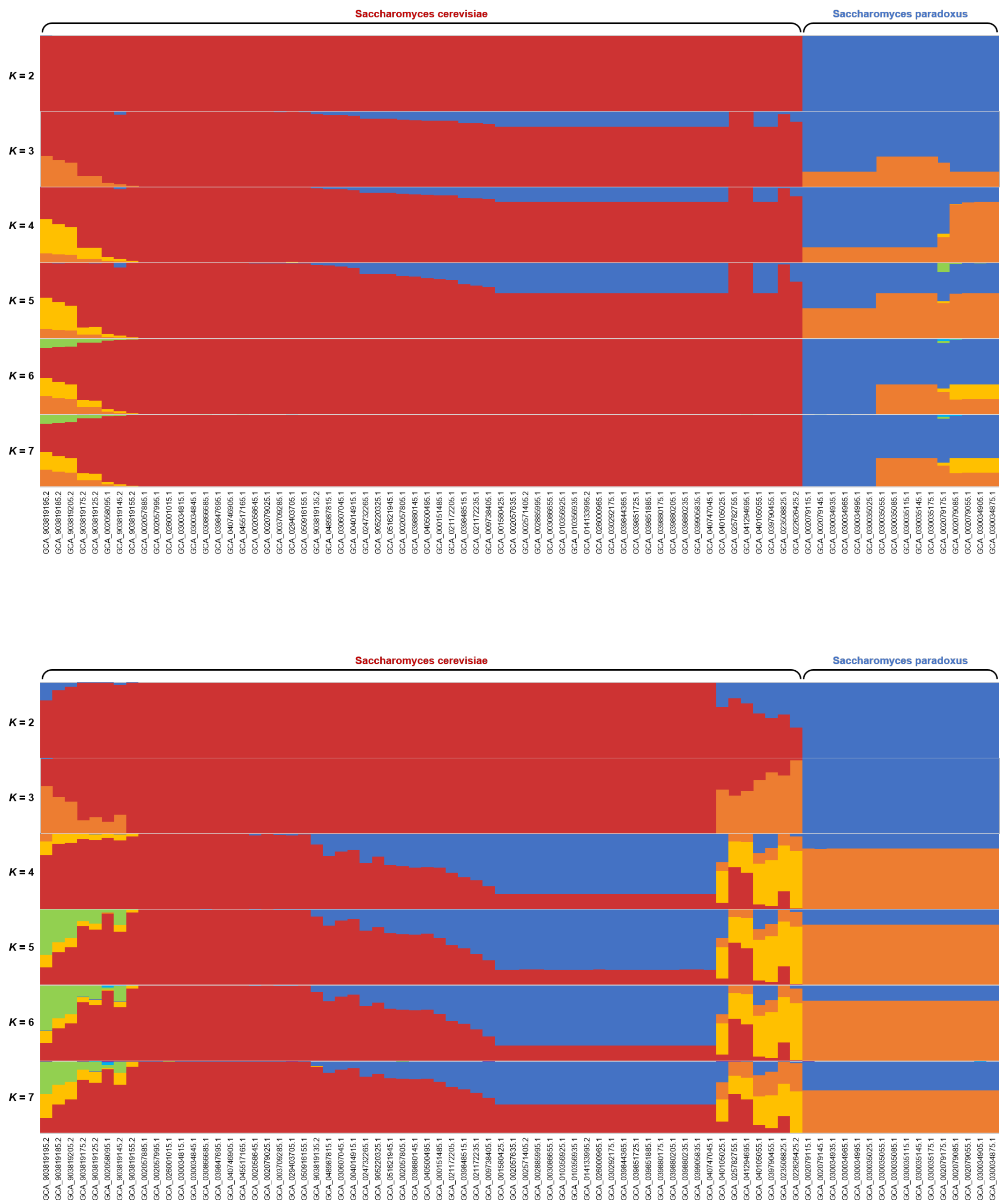


Figure 1: population structure

**A**

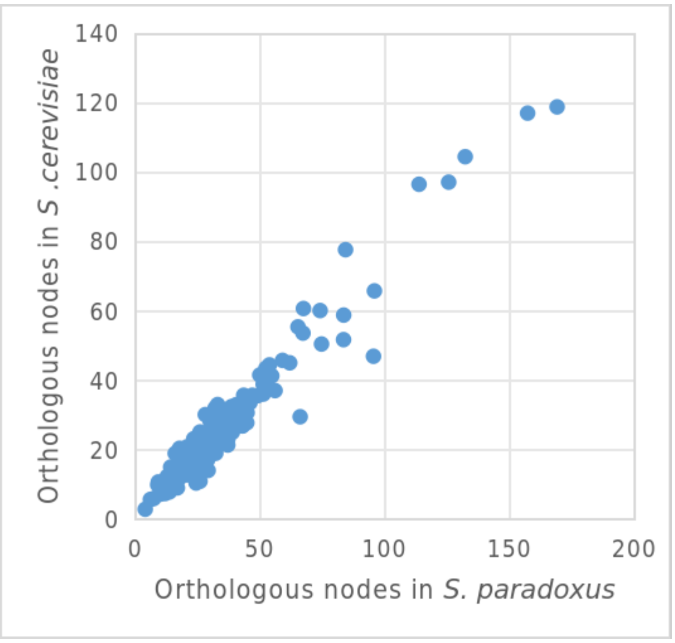


**B**

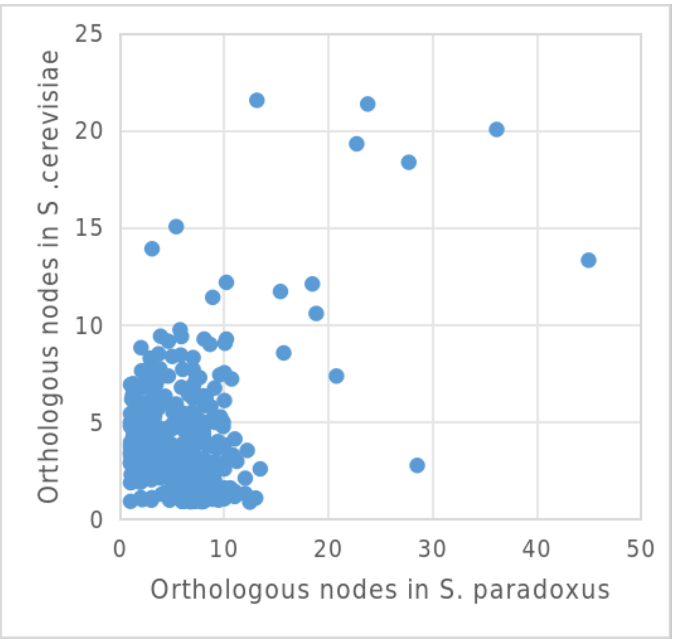


Figure 2: topoligical feature between the two yeast species

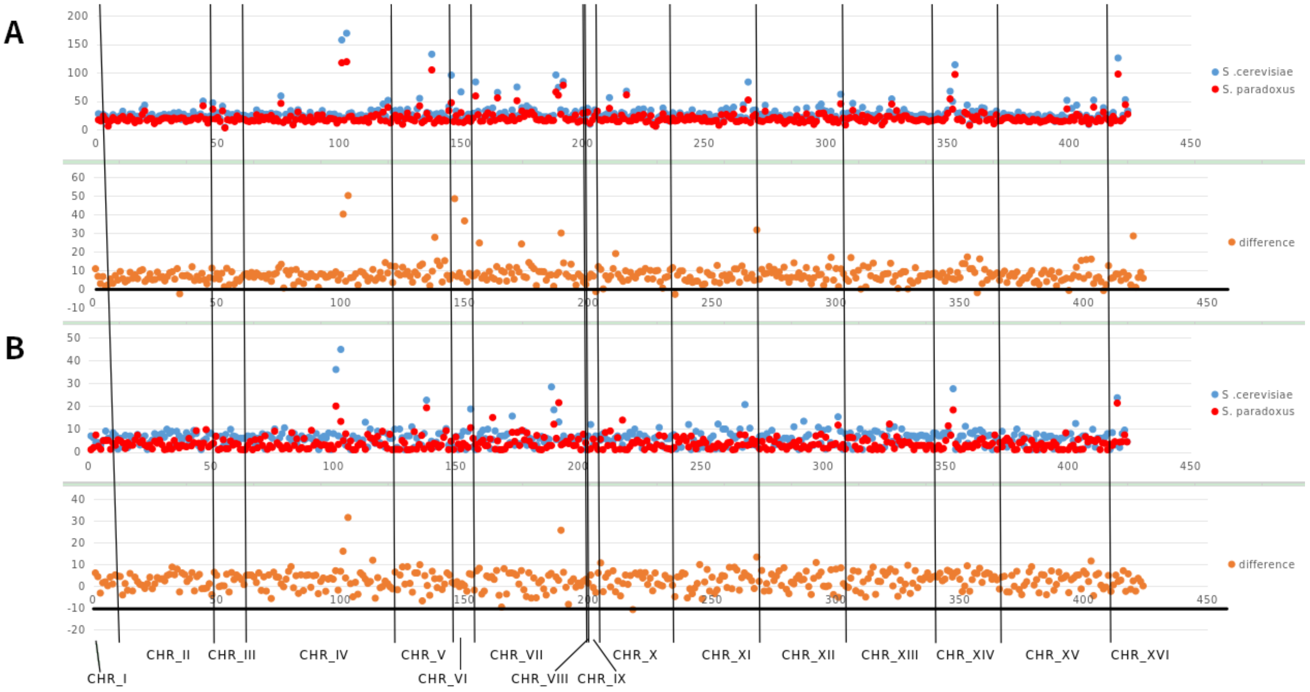


Figure 3: topoligical feature along chromosomes
